# Biological Pathways and Gene Networks Link Inflammation and Vascular Remodeling to Both Heart Failure with Preserved and Reduced Ejection Fraction in Women across Ethnicities

**DOI:** 10.1101/726208

**Authors:** Qing Liu, Kei Hang K. Chan, Alan R. Morrison, Stephen T. McGarvey, Xi Luo, James G. Wilson, Adolfo Correa, Alexander P. Reiner, Jie Li, Simin Liu, Wen-Chih Wu

## Abstract

**Introduction:** Heart failure (HF) is understudied among women; especially, genomic evidence implicating shared or unique mechanisms of HF with respect to reduced or preserved ejection fraction (HFrEF, HFpEF) is lacking across ethnic populations of women. Prior genome-wide association studies (GWAS) have identified approximately 30 suggestive genetic variants for HF, although none have been specifically linked to HFrEF or HFpEF.

**Objectives:** We aimed to define, replicate, and annotate genetic variants to HFrEF, HFpEF, or both, as well as to investigate potential biological mechanisms underlying HFrEF and HFpEF among African American (AA) and European American (EA) women in three well-characterized, high-quality prospective cohorts, the Women’s Health Initiative (WHI) study, the Jackson Heart Study (JHS), and the Framingham Heart Study (FHS).

**Methods:** GWAS analysis on HFrEF and HFpEF were first performed among 7,982 AA and 4,133 EA in the WHI, followed by pathway analysis employing two independent methodological platforms (GSA-SNP and Mergeomics) curating KEGG, Reactome, and BioCarta pathway databases. GWAS signals and biological pathways identified using the WHI were replicated in the JHS and FHS. For all replicated pathways, we performed cross-phenotype and cross-ethnicity validation analyses to examine shared pathways between HFrEF and HFpEF, and phenotype-specific pathways, across ethnicities. We further prioritized key driver genes for HF according to specific pathways identified.

**Results:** We validated one previously reported genetic locus and identified six new ones, among which one locus was allocated to HFrEF and five to HFpEF. Additionally, we defined five biological pathways shared between HFrEF and HFpEF and discovered six HFpEF-specific pathways. These pathways overlapped in two main domains for molecular signaling: 1) inflammation and 2) vascular remodeling (including angiogenesis and vascular patterning), involving key driver genes from collagen and HLA gene families.

**Conclusions:** Our network analysis of three large prospective cohorts of women in the United States defined several novel loci for HF and its subtypes. In particular, several key driver genes reinforce the mechanistic role of inflammation and vascular remodeling in the development of HF, especially HFpEF. Given that therapeutic strategies developed for left ventricular dysfunction have had limited success for HFpEF, several new targets and pathways identified and validated in this study should be further assessed in risk stratification as well as the design of potential new HF interventions.

## Introduction

According to the American Heart Association (AHA), approximately 6.5 million U.S. adults have heart failure (HF) in 2018^1^, representing a major cause of morbidity and mortality in the United States. Heart failure is phenotypically and genetically heterogeneous, and much remains unknown about the etiology for subtypes of HF, including HF with preserved ejection fraction (HFpEF) and HF with reduced ejection fraction (HFrEF). Of note as many as 40-71% of patients have HFpEF for which limited clinical treatment options are available with little to no impact on outcomes^2^. Moreover, HF is understudied in women, who experience a higher mortality than men^1^. African American (AA) women have the highest HF incidence, followed by Hispanic Americans (HA) and European Americans (EA), and morbidity and mortality are twice as high for AA women relative to EA^3-6^. Observational studies have shown that while patients with HFpEF and HFrEF often display similar clinical symptoms, they often have markedly different risk factors, underlying pathophysiological processes, and responses to clinical therapies^7^. Many therapies with unequivocal benefit in HFrEF have failed to show efficacy for HFpEF^8,9^. Thus, there is a urgent need to further research biological pathways and gene networks for HFpEF and HFrEF that could provide mechanistic insight into the disease processes and identify potential targets for novel treatment modalities, particularly for women, who are disproportionately affected by HF^1^.

Recent candidate gene studies and genome-wide association studies (GWAS) have identified several genetic loci (such as *ADRB1, USP3, ITPK1*, and *BAG3* genes^10,11^ associated with HF risk. However, these studies are primarily focused on genes associated with inherited HFrEF and are limited by reproducibility, effect size, and lack of ethnic diversity^10,11^. Although women are at higher risk of HF, no studies to date have examined/observed sex-specific genetic variants. Moreover, few studies have directly investigated the genetic mechanisms underlying the two HF subtypes, HFpEF and HFrEF^12,13^; and no studies were conducted in an ethnically diverse population of women. As genes tend to behave conjointly on HF processes, analyzing a cluster of genes with related biological functions, using an integrative pathway and network approach^14,15^, improves the statistical power to identify genetic variants of biological importance. To enhance the understanding of biological mechanisms underlying different HF phenotypes (HFpEF versus HFrEF) as well as genomic and ethnic diversity in women with HF, we therefore investigated genetic risk factors and biological pathways predisposing to HF and its subtypes among AA and EA women in the Women’s Health Initiative (WHI) study. We replicated our findings using AA women from the JHS, EA from FHS, and HA women from the WHI.

## Methods

### Study Population

#### Discovery Population

The WHI study enrolled 161,808 postmenopausal women aged between 50 and 79 years old from 1993 to 1998. The original WHI study has two major components: a partial factorial randomized clinical trial (CT) including 68,132 participants and an observational study (OS) of 93,676 participants. The detailed study design has been reported elsewhere^16^. Briefly, medical records from enrollment through September 2014 for 44,174 WHI participants, including all women randomized to the hormone trial component (n = 27,347) and all AA participants (n = 11,880) and HA participants (n = 4,947) from the CT and the OS, were sent to the University of North Carolina (UNC) for HF adjudications.

Of the participants enrolled in the WHI-OS, 8,515 self-identified AA women had consented to and were eligible for the WHI-SNP Health Association Resource (SHARe), and of the participants enrolled in the WHI hormone trial, 4,909 EA women were included in the WHI-Genomics and Randomized Trials Network (GARNET). After quality control, the standard GWAS and pathway analyses were conducted among 8,298 AA and 4,257 EA participants of the WHI. Considering the lack of replication cohort of HA participants in the WHI-SHARe, we treated the HA participants as a replication sample in the pathway analysis.

#### Population for validation and replication

The current study included three populations as validation and replication: AA in the Jackson Heart Study (JHS), EA in the Framingham Heart Study (FHS), and HA in the WHI-SHARe. Considering the WHI only enrolled postmenopausal women, we replicated the proposed analyses among women in the JHS and FHS. The main FHS enrolled three generations: the original generation (started in 1948), offspring generation (started in 1971), and generation three (started in 2005). Because of the poor measurement of left ventricular ejection fraction (LVEF) among the original generation and the relatively young age of generation three (baseline age < 40 years old), we only included the offspring generation in the analysis. In total, we conducted the proposed analyses among 1,871 AA women in the JHS, 1,764 EA women in the FHS, and 3,461 HA women in the WHI-SHARe.

### Definition of Heart Failure

In the WHI, HF adjudication was based on the Atherosclerosis Risk in Communities (ARIC) classification guidelines^17^, in which HF was defined as having acute decompensated HF (ADHF) and chronic stable HF^18^. Participants with adjudicated HF were further classified as HFrEF or HFpEF according to their LVEF. For patients with ADHF, those with LVEF < 45% were considered as HFrEF and those with LVEF ≥ 45% were considered as HFpEF. For patients with chronic stable HF, baseline LVEF or lowest estimated LVEF on medical records were used to classify HF subtypes. Similar criteria were applied to replication cohorts, the JHS and FHS. Participants without LVEF were excluded from the analysis (n=440).

### Genotype Data

Genome-wide genotyping of the WHI-SHARe and JHS participants were performed using the Affymetrix 6.0 array (Affymetrix, Inc, Santa Clara, CA), and WHI-GARNET and FHS participants were genotyped using Illumina HumanOmni1-Quad SNP platform (Illumina, Inc, San Diego, CA). As the gene chips used for genotyping are designed to capture common genetic variants, genetic variants with frequency ≥ 5% were genotyped. Reference panels from the 1000 Genomes (1000G) Project Consortium (Version 3, March 2012 release), which provide near complete coverage of common genetic variation with minor allele frequency ≥ 0.5%, were used for genotype imputation.

### Statistical Analysis

#### Genome-wide association analyses

We performed standard GWAS analysis for HFrEF and HFpEF for AA and EA women, using multivariable logistic regressions. The regression models were implemented using allelic dosage at each SNP (single-nucleotide polymorphism) as the independent variable, with covariate adjustment for age, age^2^, and first four principal components (PCs) for global ancestry in all three cohorts. We also adjusted for region in two WHI cohorts, and randomized hormone treatment group and baseline hysterectomy status in the WHI-GARNET study. Since the associations between germline genetic variants and HF are not confounded by demographic and lifestyle factors, no other confounders were adjusted in the GWAS analysis. The general form of the GWAS model is specified as follows:

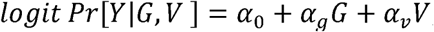

where Y denotes HF subtype, G denotes SNPs, and V denotes adjusted covariates. Common genetic variants reaching the suggestive significance (5× 10^−6^) were identified as potential GWAS hits. Suggestive SNPs were validated in the JHS and FHS using nominal *P* value < 0.05, followed by cross-ethnicity meta-analyses combining AA, EA, and HA women from WHI, JHS, and FHS using METAL^19^ (FDR-adjusted q value < 0.05).

#### Pathway analysis

We obtained knowledge-driven metabolic and signaling pathways from three databases: the Kyoto Encyclopedia of Genes and Genomes (KEGG)^20^, Reactome^21^, and BioCarta^22^. SNPs showing potential associations with HF subtypes (*P* value <0.05 in GWAS) were mapped to relevant genes based on their chromosome locations or functions, and further mapped to biological pathways. Each pathway was tested for enrichment of genetic signals for HFrEF and HFpEF by ethnic groups. To avoid potential biases due to a particular method, we applied two different well-established methods based on known biological pathways: 1) GSA-SNP^14^, and 2) Mergeomics^15^. Pathways were defined as significant if they met the following criteria: 1) identified by both methods from the WHI study with a FDR-adjusted q value < 0.2; and 2) validated by GSA-SNP or Mergeomics with a significant *P* value after Bonferroni correction in JHS (as replication of WHI-AA) and FHS (as replication of WHI-EA). We then performed cross-phenotype and cross-ethnicity analyses in WHI to examine shared pathways between HFrEF and HFpEF, as well as phenotype-specific pathways, across ethnicities (AA, EA, and HA).

#### Key Driver Analysis for Identification of Key Regulatory Genes for HF-related Pathways

As hundreds of genes are involved in the biological pathways, we seek to further prioritize key driver (KD) genes, defined as genes that played a central role in the disease progress and once perturbed, should have major impact on many other genes. We integrated all genes involved in significant pathways with seven Bayesian networks and one protein-protein interaction network using KD analysis methods^15,23,24^. We designed a normalized rank score (NRS) to summarize the consistency and strength of identified KD genes across multiple networks^25^, where 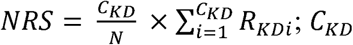 is the count of networks from which a KD was identified; *C*_*KD*_ is then normalized by total number of networks N to represent the consistency of a KD across all networks tested (Bayesian networks from seven tissues, including adipose, blood, brain, islet, liver, kidney, and muscle, and one protein-protein interaction). The KD strength is represented by the summation of normalized statistical rank in each network *i* (*R*_*KDi*_) across all networks from which the KD is identified; 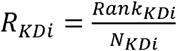, which was calculated by dividing the rank of a KD based on the *P* values of the Fisher exact test in descending order (*Rank*_*KDi*_) by the total number of KDs identified from a network *i* (*N*_*KDi*_). KDs with high NRS were those with high network enrichment for pathways and high consistency across tested networks.

## Results

Among WHI participants, 860 (10.4%) AA, 601 (14.1%) EA, and 165 (4.7%) HA were initially identified as having HF. After excluding those without LVEF measurement (316 WHI-AA and 124 WHI-EA), we performed primary analyses among 7,982 AA and 4,133 EA women in the WHI, and replication analyses among 1,853 AA women in the JHS, 1,755 EA women in the FHS, and 3,461 HA women in the WHI. The descriptive statistics on demographic and lifestyle factors of each study population are shown in **Table 1**. Compared to WHI-EA, WHI-AA women were younger in age and less physically active, had higher BMI and lower intakes of alcohol and total calories, and with a higher proportion of cardiovascular disease and diabetes.

**Table 1.** Baseline Characteristics of African and European American Women in Study Populations ^a^

|  | Discovery Populations |  | Replication Populations <sup>b</sup> |  |  |
| --- | --- | --- | --- | --- | --- |
|  | WHI-AA<br>(N=7,982) | WHI-EA<br>(N=4,133) | JHS-AA<br>(N=1,853) | FHS-EA<br>(N=1,755) | WHI-HA<br>(N=3,461) |
| **No. of participants with HF, n (%) <sup>c</sup>** | 544 (6.8) | 477 (11.5) | 166 (9.0) | 108 (6.2) | 100 (2.9) |
| HFpEF, n (%) | 345 (63.4) | 304 (63.7) | 140 (84.3) | 103 (95.4) | 67 (67.0) |
| HFrEF, n (%) | 199 (36.6) | 173 (36.3) | 26 (15.7) | 5 (4.6) | 33 (33.0) |
| Years of follow-up, year (SD) | 10.0 (5.2) | 8.9 (5.3) | 6.4 (2.3) | 38.9 (4.8) | 10.4 (5.3) |
| **Age, year (SD)** | 61.5 (7.0) | 65.6 (6.9) | 55.5 (12.7) | 34.8 (9.8) | 60.2 (6.7) |
| **Region, n(%) <sup>d</sup>** |  |  |  |  |  |
| Northeast | 1410 (17.7) | 1039 (25.1) | -- | -- | 436 (12.6) |
| South | 3902 (48.9) | 879 (21.3) | -- | -- | 1423 (41.1) |
| Midwest | 1854 (23.2) | 1185 (28.7) | -- | -- | 126 (3.6) |
| West | 816 (10.2) | 1030 (24.9) | -- | -- | 1476 (42.6) |
| **Current smoking, n (%)** | 901 (11.3) | 449 (10.9) | 202 (10.9) | 723 (41.2) | 233 (6.7) |
| **Cardiovascular disease, n (%) <sup>e</sup>** | 1355 (17) | 647 (15.7) | 191 (10.3) | 0 (0) | 356 (10.3) |
| **Diabetes, n (%)** | 1054 (13.2) | 285 (6.9) | 436 (23.5) | 8 (0.5) | 278 (8) |
| **BMI, kg/m<sup>2</sup> (SD)** | 31 (6.5) | 29.6 (6.1) | 33.3 (7.8) | 24 (4.4) | 28.9 (5.8) |
| **Physical Activity, MET-h/week (SD) <sup>d</sup>** | 9.7 (12.7) | 10.2 (12.8) | -- | -- | 10.8 (13.8) |
| **Alcohol drinking, serving/week (SD)** | 1.1 (3.9) | 2.3 (5.1) | 1.5 (6.2) | 3.5 (5.8) | 1.3 (3.9) |
| **Total energy intake, kcal/day (SD)** | 1614.1 (759.5) | 1683.3 (663.1) | 1826.2 (751.8) | 1757.0 (579.6) | 1656.7 (777.4) |

Abbreviations: AA (African American), ADHF (acute decompensated heart failure), BMI (body mass index), EA (European American), FHS (Framingham Heart Study), HA (Hispanic American), HF (heart failure), HFpEF (heart failure with preserved ejection fraction), HFrEF (heart failure with reduced ejection fraction), JHS (Jackson Heart Study), LVEF (left ventricular ejection fraction), SD (standard deviation), WHI (Women's Health Initiative Study).
<sup>a</sup> Continuous variables were presented as mean (SD).
<sup>b</sup> 62.0% in the JHS and 52.5% in the FHS participants were women.
<sup>c</sup> HF was defined as having ADHF or chronic stable HF, and further classified as HFrEF (LVEF < 45%) or HFpEF (LVEF ≥ 45%). For WHI patients with ADHF, LVEF closest to the diagnosis date of ADHF was used, and for patients with chronic stable HF, baseline LVEF or lowest estimated LVEF on medical records were used to classify HF subtypes. In the JHS, considering over 50% of ADHF patients were missing LVEF, baseline LVEF was used to determine HFrEF and HFpEF for those without coronary heart disease before HF, and for chronic stable HF patients, baseline LVEF was used. In the FHS, HF was defined as incident or prevalent congestive HF. Since LVEF was not measured at baseline, LVEF closest to the diagnosis date of HF was used to define HFrEF and HFpEF. Participants without LVEF measurement were excluded in the analysis.
<sup>d</sup> Results in the replication populations were not presented for variables which were not applicable (region) or measured in different scales (physical activity) .
<sup>e</sup> Cardiovascular disease was defined as self-reported coronary heart disease, heart failure, stroke, and peripheral artery disease at baseline.

### Identification of Significant Genetic Loci Using Standard GWAS Analysis

In the validation analysis of previously reported 30 loci for HF from the GWAS catalog^26^, we validated one locus and further allocated it to HFpEF in the WHI, JHS, and FHS populations with FDR-adjusted q value < 0.05. The validated SNP rs4420638 is located on chromosome 19 and close to *APOE* and *APOC* genes. Detailed information regarding the validated locus can be found in **Supplemental Table 1**.

The standard GWAS results for HFrEF and HFpEF within WHI-AA (n=7,982) and WHI-EA (n=4,133) are shown in the Manhattan plots (**Supplementary Figure 1**). Among AA, this discovery analysis revealed one significant (*P* < 5×10^−8^, rs35900865) and 57 suggestive (*P* < 5×10^−6^) SNPs related to HFrEF, and three significant (*P* < 5×10^−8^, rs7834398, rs78668964, and rs12203350) and 94 suggestive (*P* < 5×10^−6^) SNPs related to HFpEF. Among EA, we failed to identify significant SNPs, but found 50 suggestive (*P* < 5×10^−6^) SNPs related to HFrEF and 47 SNPs for HFpEF.

In the replication analysis for AA women among JHS (n=1,853) participants, eight SNPs from four loci (lead SNPs: rs12067046, rs114553497, rs10229703, and rs149663839) out of 94 SNPs for HFpEF reached the threshold of *P* < 0.05. In the replication analysis for EA women among FHS (n=1,755) participants, one SNP (rs12719020) reached the *P* < 0.05 threshold among the 50 suggestive SNPs for HFrEF; and 19 SNPs (concentrated on chromosome 16, lead SNP: rs12599260) among the suggestive 47 SNPs for HFpEF reached the threshold of *P* < 0.05. The effect of all lead SNPs on HF was in the same direction in the discovery population and the ethnicity-specific replication population. In the cross-ethnicity meta-analysis combining AA, EA, and HA women from the WHI, JHS and FHS, all loci passed ethnicity-specific validation and were further validated with FDR-adjust q value of < 0.05 (**Table 2**). More information regarding the newly discovered loci can be found in **Supplemental Table 1**.

**Table 2.**
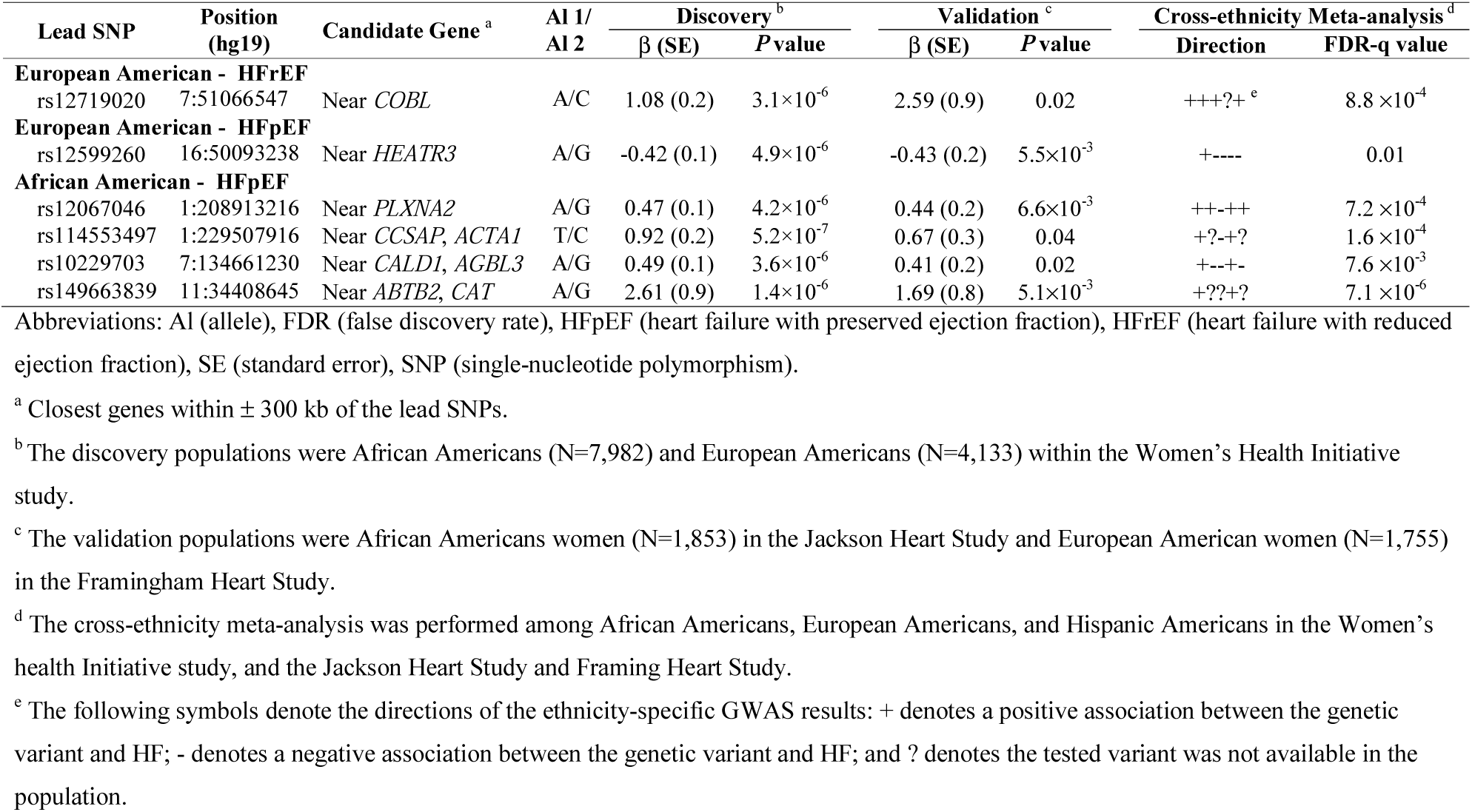
Newly Discovered Loci for Heart Failure among Women across Ethnicities in the Women’s Health Initiative Study, Jackson Heart Study and Framingham Heart Study.

### Identification of Biological Pathways Using Integrative Pathway Analysis

We initially identified 21 pathways for HFrEF (nine for EA and 12 for AA) and 42 pathways for HFpEF (31 for EA and 17 for AA) among WHI participants, of which 11 pathways were validated for HFrEF and 15 pathways for HFpEF, among the JHS and FHS women. The results of cross-phenotype and cross-ethnicity analysis were presented in **Table 3 and Supplemental Tables 2 and 3**. Based on the functions of the pathways, we identified two main overarching domains with some cell signaling and metabolism common to both: 1) angiogenesis and vascular patterning and 2) inflammation. Five pathways, emerging from angiogenesis and vascular patterning, were shared between HFrEF and HFpEF across AA, EA, and HA women, namely, extracellular matrix (ECM)-receptor interaction, cell adhesion molecules (CAMs), axon guidance, netrin-1 signaling, and developmental biology (**Figure 1)**. The five shared pathways were highly interconnected as demonstrated by a shared common set of 256 genes among them (**Figure 2**).

**Figure 1.**
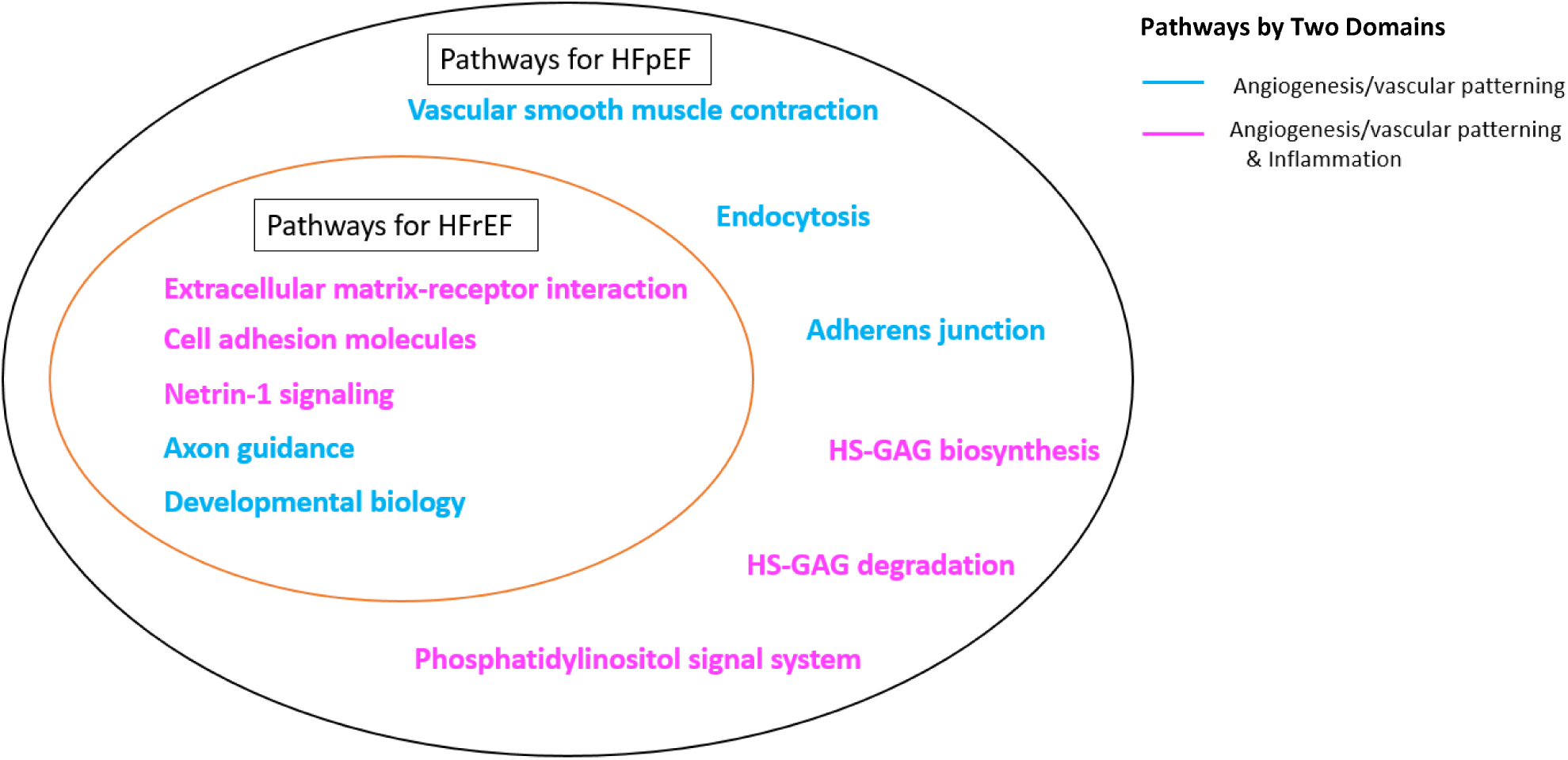
Venn Diagram for Biological Pathways Enriched for HFrEF and HFpEF among African and European American Women across Ethnicities. Abbreviations: GAG (glycosaminoglycan), HFpEF (heart failure with preserved ejection fraction), HFrEF (heart failure with reduced ejection fraction), HS (Heparan sulfate).

**Figure 2.**
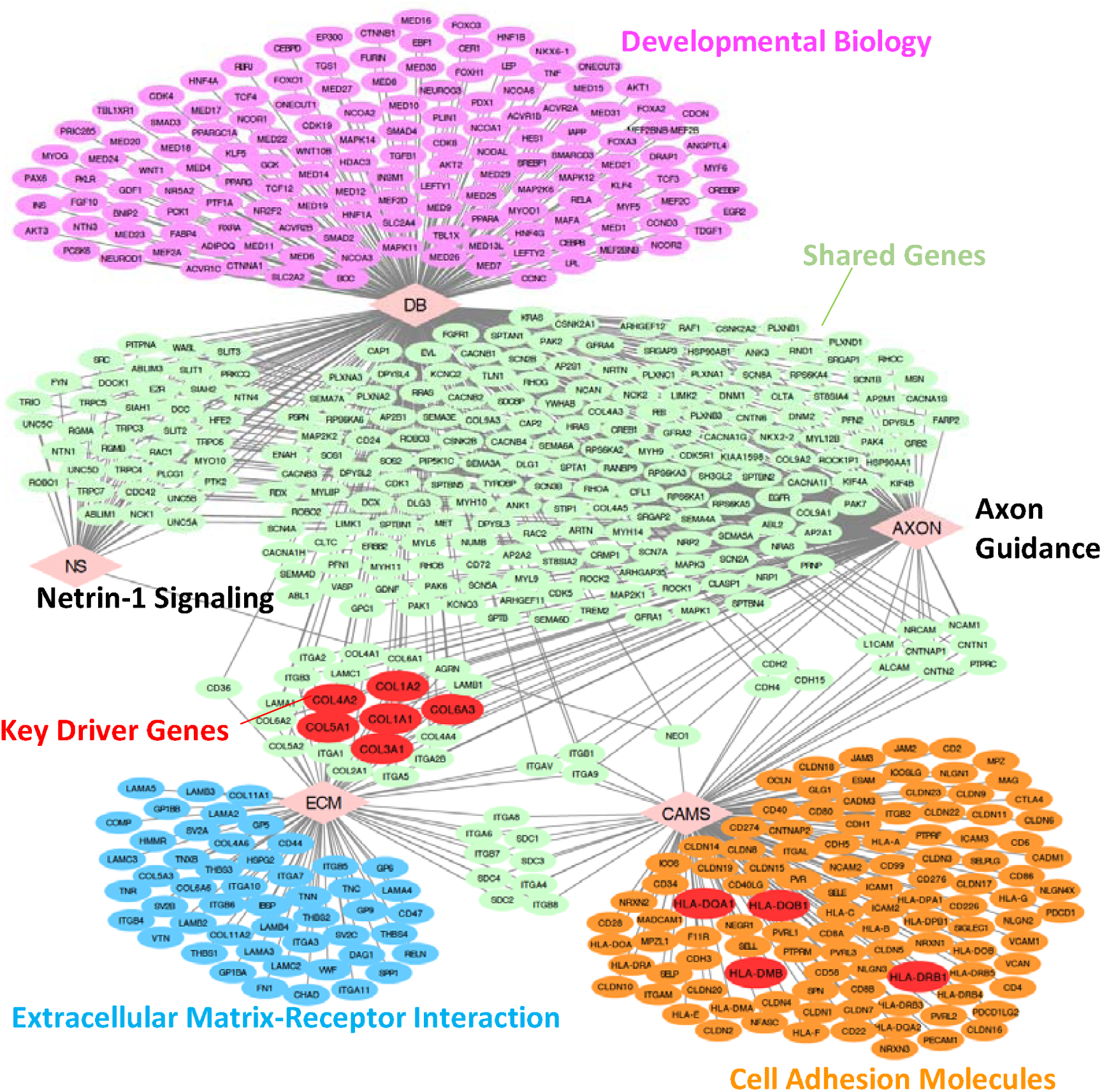
Network of 5 Pathways Enriched for HFrEF and HFpEF with Top 10 Key Driver Genes among African and European American Women. The diamond nodes represent pathway and the ellipse modes represent genes, and the edge shows the interaction, that is, the association between a gene and a pathway. The color nodes are: red, top 10 key driver genes; light green, genes involved in ≥ 2 pathways; others are pathway-specific genes. The figure was created using Cytoscape^86^. Abbreviations: AXON (axon guidance), CAMS (Cell adhesion molecules), DB (developmental biology), ECM (Extracellular matrix-receptor interaction), HFpEF (heart failure with preserved ejection fraction), HFrEF (heart failure with reduced ejection fraction), NS (Netrin-1 signaling).

**Table 3.**
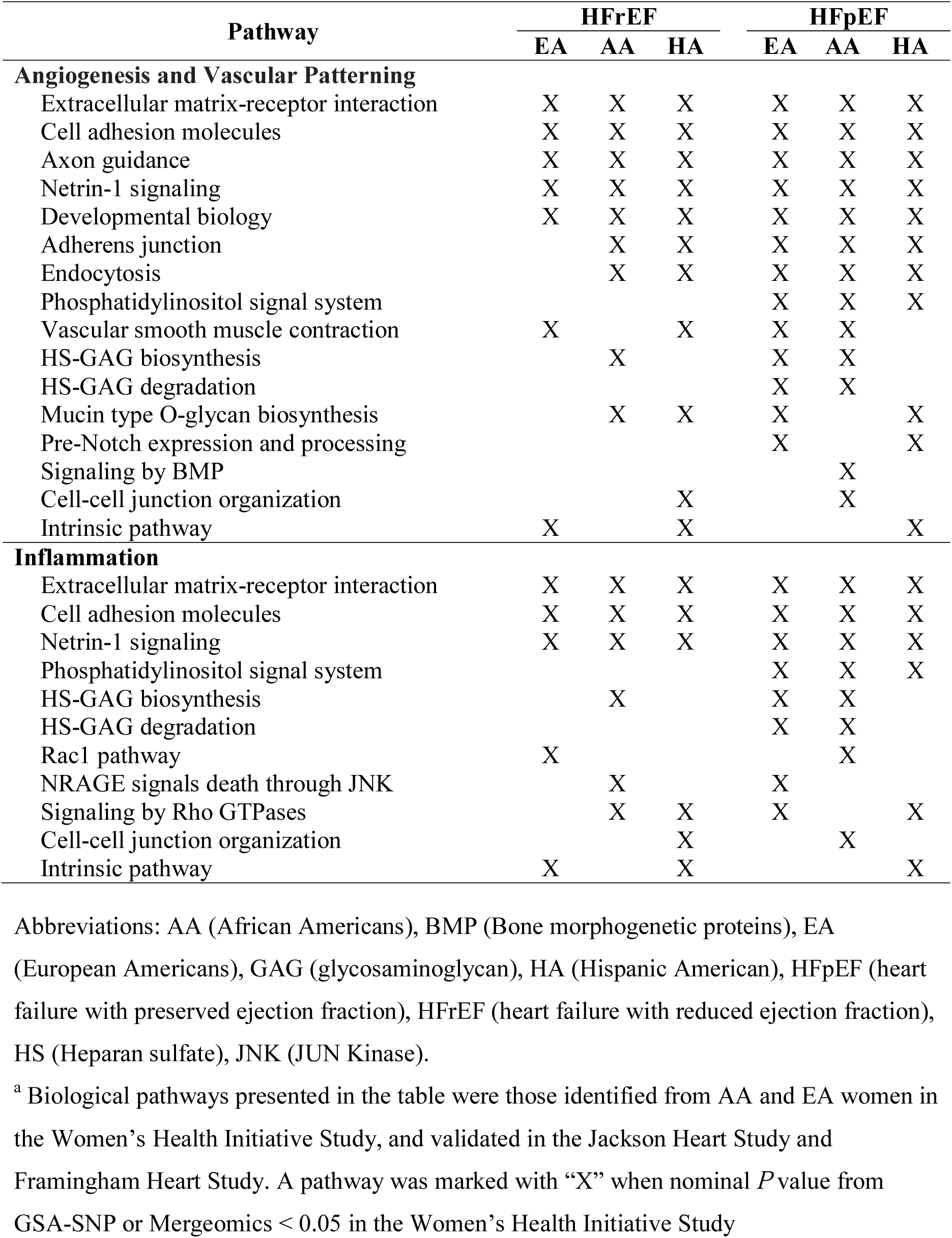
Biological Pathways Enriched for HFrEF and HFpEF among African and European American Women across Ethnicities ^a^.

| Pathway | HFrEF |  |  | HFpEF |  |  |
| --- | --- | --- | --- | --- | --- | --- |
|  | EA | AA | HA | EA | AA | HA |
| **Angiogenesis and Vascular Patterning** |  |  |  |  |  |  |
| Extracellular matrix-receptor interaction | X | X | X | X | X | X |
| Cell adhesion molecules | X | X | X | X | X | X |
| Axon guidance | X | X | X | X | X | X |
| Netrin-1 signaling | X | X | X | X | X | X |
| Developmental biology | X | X | X | X | X | X |
| Adherens junction |  | X | X | X | X | X |
| Endocytosis |  | X | X | X | X | X |
| Phosphatidylinositol signal system |  |  |  | X | X | X |
| Vascular smooth muscle contraction | X |  | X | X | X |  |
| HS-GAG biosynthesis |  | X |  | X | X |  |
| HS-GAG degradation |  |  |  | X | X |  |
| Mucin type O-glycan biosynthesis |  | X | X | X |  | X |
| Pre-Notch expression and processing |  |  |  | X |  | X |
| Signaling by BMP |  |  |  |  | X |  |
| Cell-cell junction organization |  |  | X |  | X |  |
| Intrinsic pathway | X |  | X |  |  | X |
| **Inflammation** |  |  |  |  |  |  |
| Extracellular matrix-receptor interaction | X | X | X | X | X | X |
| Cell adhesion molecules | X | X | X | X | X | X |
| Netrin-1 signaling | X | X | X | X | X | X |
| Phosphatidylinositol signal system |  |  |  | X | X | X |
| HS-GAG biosynthesis |  | X |  | X | X |  |
| HS-GAG degradation |  |  |  | X | X |  |
| Rac1 pathway | X |  |  |  | X |  |
| NRAGE signals death through JNK |  | X |  | X |  |  |
| Signaling by Rho GTPases |  | X | X | X |  | X |
| Cell-cell junction organization |  |  | X |  | X |  |
| Intrinsic pathway | X |  | X |  |  | X |

Abbreviations: AA (African Americans), BMP (Bone morphogenetic proteins), EA (European Americans), GAG (glycosaminoglycan), HA (Hispanic American), HFpEF (heart failure with preserved ejection fraction), HFrEF (heart failure with reduced ejection fraction), HS (Heparan sulfate), JNK (JUN Kinase).
<sup>a</sup> Biological pathways presented in the table were those identified from AA and EA women in the Women's Health Initiative Study, and validated in the Jackson Heart Study and Framingham Heart Study. A pathway was marked with "X" when nominal *P* value from GSA-SNP or Mergeomics < 0.05 in the Women's Health Initiative Study

In addition, we found six pathways specifically enriched for HFpEF across AA and EA, namely, adherens junction, endocytosis, phosphatidylinositol signal system, vascular smooth muscle contraction, and heparan sulfate/heparin (HS)-glycosaminoglycan (GAG) biosynthesis and degradation; all of which corresponded to the domain of angiogenesis and vascular patterning (**Figures 1 and 3**). Of the aforementioned six HFpEF-specific pathways, the first three pathways were further replicated in the WHI-HA participants.

**Figure 3.**
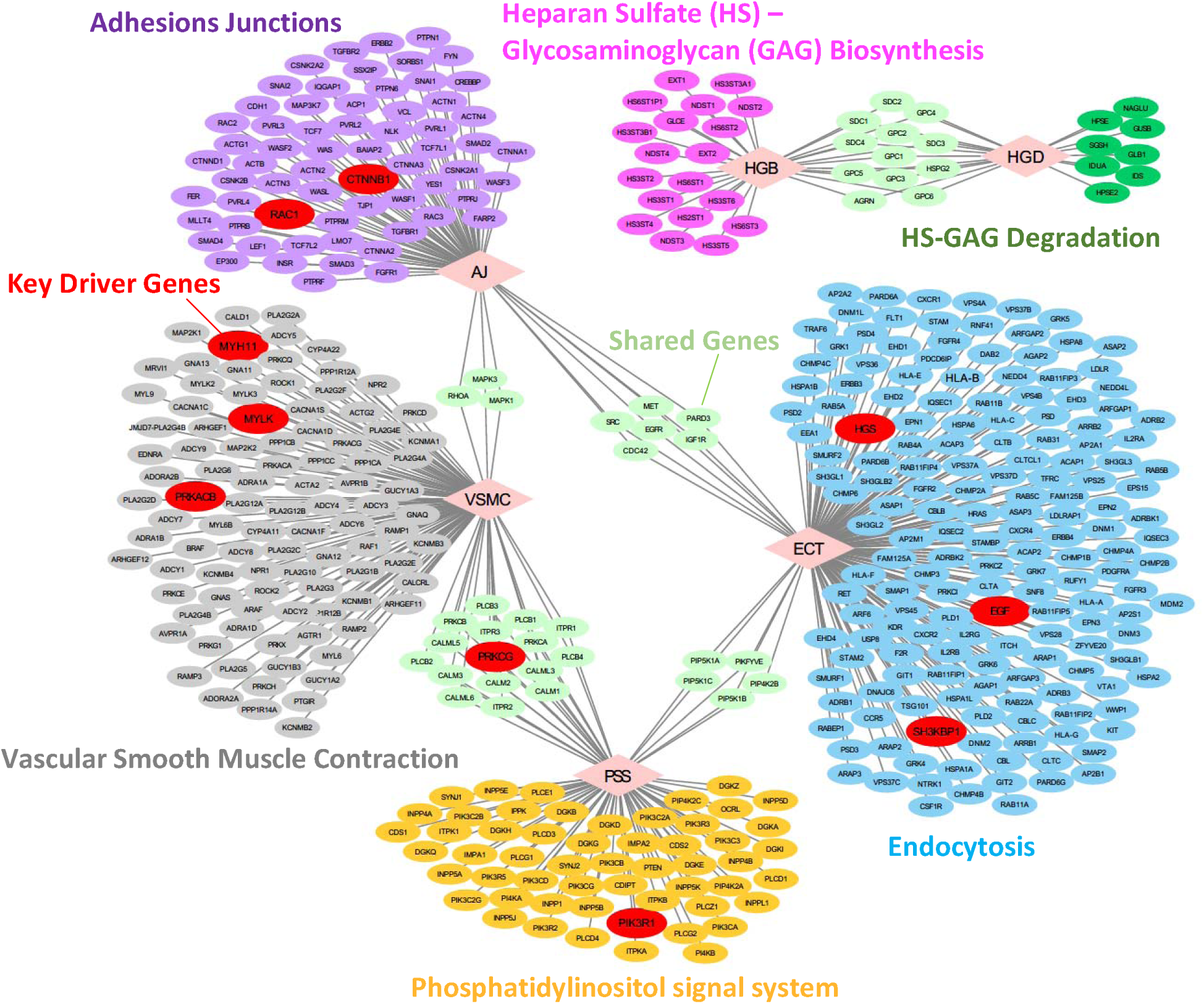
Network of 6 Pathways Enriched for HFpEF with Top 10 Key Driver Genes among African and European American Women. The diamond nodes represent pathway and the ellipse modes represent genes, and the edge shows the interaction, that is, the association between a gene and a pathway. The color nodes are: red, top 10 key driver genes; light green, genes involved in ≥ 2 pathways; others are pathway-specific genes. The figure was created using Cytoscape^86^. Abbreviations: AJ (adherens junction), ECT (endocytosis), HGB (heparan sulfate-glycosaminoglycan biosynthesis), HGD (heparan sulfate-glycosaminoglycan degradation), HFpEF (heart failure with preserved ejection fraction), PSS (phosphatidylinositol signal system), VSMC (vascular smooth muscle contraction).

### Identification of Key Drivers for HFpEF and HFrEF

In the KD analysis to identify potential genes that played a central role in the significant pathways for HF, we used eight different regulatory or interaction networks that capture gene-gene or protein-protein interactions among the pathways. The top 10 KD genes for the five shared pathways (developmental biology, axon guidance, netrin-1 signaling, ECM-receptor interaction, and CAMs) between HFrEF and HFpEF across two ethnicities are *COL1A1, COL1A2, COL3A1, COL4A2, COL5A1*, and *COL6A3* from ECM and axon guidance pathways, and *HLA-DQA1, HLA-DQB1, HLA-DRB1*, and *HLA-DMB* from CAMs pathway (**Figure 2 and Supplemental Figure 2**). For the six pathways specific for HFpEF, the top 10 KD genes are *MYH11, MYLK* and *PRKACB* from vascular smooth muscle contraction, *PRKCG, PIK3R1* from phosphatidylinositol signal system, *HGS, EGF*, and *SH3KBP1* from endocytosis, and *CTNNB1* and *RAC1* from adherens junction (**Figure 3 and Supplemental Figure 2**). Variants in the identified top KD genes collectively account for 15-19% and 15-16% variations of HFrEF and HFpEF among women in the WHI.

## Discussion

In this GWAS analysis of 7,982 AA and 4,133 EA women from the WHI, we validated one previously reported genetic locus and allocated it to HFpEF, and additionally discovered one HFrEF and five HFpEF novel genetic loci of potential importance. Also, five biological pathways appeared to be shared for both HFrEF and HFpEF across AA, EA, and HA women, and six pathways were specific for HFpEF across AA and EA women. Our results suggested the presence of core mechanisms across HF subtypes (HFrEF and HFpEF), such as vascular remodeling and inflammation alone with some common overlapping mechanisms of cell signaling and metabolism. It is important to note the paucity of cardiomyocyte-specific gene variants, including those for nuclear envelope proteins, sarcomere proteins, cytoskeletal, and calcium regulatory proteins, given their extensive involvement in familial dilated cardiomyopathies. Our data highlight the genetic and/or biological significance of the vascular remodeling and inflammation, rather than that of the cardiomyocyte in the acquisition of HFpEF and HFrEF.

Given the increased diversity of gene involvement, the genetic architecture underlying HFrEF and HFpEF remains challenging to delineate. We validated one previously reported locus close to *APOE* and *APOC*, and further allocated it to HFpEF. Genes in the apolipoprotein family (*APOE, APOC1, APOC2*, etc.) encode lipid transport proteins that regulate cholesterol metabolism and are associated with obesity and cardiovascular disease^27,28^. Of note, we were not able to validate other suggestive HF loci reported from previous European-based cohorts, which may be due to effect modifications by sex and/or ethnicity.

In addition, we discovered one HFrEF and five HFpEF loci from intergenic regions. Variant rs12719020, associated with HFrEF, is located upstream (< 20 Kb) to *COBL*, a gene related to vasculitis and type 1 diabetes^29^. For variants associated with HFpEF, rs12067046 is located 500 Kb downstream of *PLXNA2*, which is related to the development of blood vessel^30^ and inflammatory-induced immune disorders^31^; the linkage disequilibrium (LD) block around rs12599260 is upstream (5 Kb) to *HEATR3*, which regulates inflammatory immune response^32^; rs149663839 is located upstream (50 Kb) to *CAT*, a key antioxidant enzyme, which is hypothesized to play a role in the development of many chronic or late-onset diseases such as HF^33^; *ACTA1* (60 Kb to the LD block around rs114553497) and *CALD1* (5 Kb to rs10229703) are fundamental genes for skeletal/smooth muscle contraction and had been linked to pulmonary hypertension in animal studies^34,35^ (**Table 2 and Supplemental Table 1**).

Genetic pathway and network analysis, as novel approaches to integrate genetic signals that complements current GWAS analysis, have been yielded new insight into the biology of coronary heart disease^36^, type 2 diabetes^25^, obesity^37^, and LV function^38^. Our pathway-based analysis revealed five consistent pathways between HFrEF and HFpEF across the two ethnicities (**Table 3)**. All the five pathways were linked to angiogenesis and vascular patterning, among which three pathways, ECM-receptor interaction, CAMs, and Netrin-1 signaling were also linked to inflammation (**Figure 1**). From the five pathways shared by both HFrEF and HFpEF, three pathways, axon guidance, ECM-receptor interaction, and CAMs, had been implicated previously in thromboembolic cardiovascular disease, type 2 diabetes, and LV function^25,38^.

There appears to be a substantial overlap in the major areas of inflammation and angiogenesis among shared pathways between HFrEF and HFpEF. The ECM is an intricate network composed of multidomain macromolecules organized to support mechanical and structural properties of cells and tissue but also to control behavioral characteristics of cells, including proliferation, adhesion, migration, polarity and differentiation^39,40^. Major components include collagens, proteoglycans, elastin, and cell-binding glycoproteins, each with distinct physical and biochemical properties. ECM molecules connect to the cells through integrins, syndecans, and other receptors which provides signaling input in addition to mechanical support^41^. This ECM-receptor interaction contributes to angiogenesis and vascular patterning in multiple ways, including the organization and maintenance of gradients for angiogenic factors like vascular endothelial growth factor (VEGF)-A^42,43^. Endothelial ECM receptors like intergrins play a critical role in adhesion and migration via control of cytoskeletal dynamics while at the same time directing cell-cell interactions through pathways like Notch signaling in order to coordinate sprouting and tube organization in early capillary networks^44^. ECM-CAM interactions also have the ability to influence inflammatory state of both vascular and immune cells through focal adhesion complexes comprised of integrins, protein kinases such as focal adhesion kinase (FAK), Src and many other kinases, adaptor proteins such as Shc, signaling intermediates such as phosphoinositide 3-kinase (PI3K), Rho and Rac GTPases, and actin binding cytoskeletal proteins^45^. Further, cardiac ECM (primarily collagen I) may also play a critical role in providing a platform for cardiomyocytes to maintain structure and function, and any change in ECM properties following an insult has potential to drive the progression toward HF, including myocardial fibrosis and altered ECM protein orientation^46,47^. Details regarding the main functions of other identified pathways can be found in **Supplemental Table 4.**

Taken together, the above identified pathways highlight the previously known but likely underappreciated importance of angiogenesis and vascular patterning as well as inflammation on HF that appear to link HFrEF and HFpEF to other cardio-metabolic health outcomes via multiple mechanisms. The fact that these pathways were consistently identified across multiple ethnicities further highlights a convergent or central role in the joint mechanisms between interrelated cardiac diseases.

We additionally identified six pathways specifically for HFpEF across ethnicities. All the six pathways were linked to angiogenesis and vascular patterning, from which three pathways (phosphatidylinositol signal system, HS-GAG biosynthesis and degradation) were additionally linked to inflammation (**Figure 1**). None of the six HFpEF pathways have previously been implicated in pathway-based studies, making these findings novel.

Specifically, the phosphatidylinositol signaling system is critically linked to inflammation and vascular remodeling, and in particular angiogenesis and vascular patterning. Activation of the PI3K pathway can occur in response to a variety of extracellular (e.g., ECM, CAM) as well as growth factor (e.g., fibroblast growth factor, VEGF-A) signaling and can regulate broad spectrum of molecular functions which involve proliferation, adhesion, migration, invasion, metabolism and cell survival^48,49^. Activation of the PI3K pathway involves recruitment via Src-homology 2 (SH2) to phosphotyrosine residues on the intracellular portion of membrane receptors, followed by phosphorylation of phosphatidylinositol-4,5-bisphosphate (PIP2) to generate the second messenger molecule PIP3. The Akt family of serine/threonine kinases has been shown to be the primary downstream mediator of the effects of PI3K. Through the phosphorylation of IκB kinase (IKK) and activation of nuclear factor κ B (NF-κB) transcriptional activity, Akt leads to upregulation of inflammatory and prosurvival genes^50-53^. Akt can also activate mTOR, resulting in stabilization of hypoxia inducible factor-1 (HIF-1) and consequent expression of VEGF-A in order to promote angiogenesis^54-56^. PI3K signaling may also play a role in regulating cardiomyocyte size, survival, and inflammation during cardiac hypertrophy and HF, in part via calcium signaling^57,58^. Details regarding the main functions of other identified pathways can be found in **Supplemental Table 4.** Detailed knowledge of these relationships at the molecular level will allow researcher to understand the distinct mechanisms underlying HFpEF and enable the development of effective therapeutic strategies.

In addition, we observed two pathways with highly significant *P* values: Rac1 pathway for HFrEF within EA and HFpEF within AA, and signaling by bone morphogenetic proteins (BMP) pathway for HFpEF among AA (**Supplemental Tables 2, 3, and 4**). Rac1 is a GTPase protein, a member of the Rac subfamily of the Rho family of GTPases, and plays a critical role in inflammation and vascular remodeling. Activated Rac1 can promote NF-κB signaling and reactive oxygen species (ROS) production via nicotinamide adenine dinucleotide phosphate (NADPH) oxidase, both known activators of the NACHT, LRR, and PYD domains-containing protein 3 (NLRP3) inflammasome protein complex that promotes expression of the critical inflammatory cytokine, interleukin-1β (IL-1β)^59-61^. A number of studies have found that activation of Rac1 is associated with atrial fibrillation^62^, atherosclerotic calcification^63^, cardiac hypertrophy^64^, and HF^65^, suggesting Rac1 may have strong potential as a new therapeutic target^66^. BMPs belong to transforming growth factor beta (TGFβ) superfamily, which is one of the most potent profibrogenic cytokine systems governing cardiac fibrosis^67^. BMP signaling is also increasingly recognized for its influence on endocrine-like functions in postnatal cardiovascular and metabolic homeostasis^68^. Some BMP molecules, such as BMP9 and BMP10, had been found to reduce pulmonary arterial hypertension, cardiac fibrosis, and myocardial infarction, thereby providing potentially benefits for HF patients^68,69^.

The KD gene analysis prioritized KD genes of coronary heart disease^70^, type 2 diabetes^25^, and obesity^71^, but has not been performed for HF. In our KD gene analysis based on shared pathways between HFrEF and HFpEF, we found that the KD genes belong to the collagen gene family, shared between axon guidance and ECM-receptor interaction, and HLA genes from CAMs pathway. HLA gene family members are components of the major histocompatibility complex (MHC) and play a central role in the immune system with established allelic contributions to type 1 diabetes susceptibility^72^, and a host of inflammatory disorders, including rheumatoid arthritis^73^, Sjögren’s^74^, ulcerative colitis ^75^, and systemic lupus erythematosus^76^. Collagen gene family, as previously described, encodes proteins to regulate vascular patterning and maintain the structure and function of cardiomyocytes. These KD genes further highlight the effect of angiogenesis, vascular patterning and inflammation in HF. Importantly, genes *COL1A1* and *COL3A1* were also found to be the KD genes for thromboembolic cardiovascular disease and type 2 diabetes^25^; thus, showing potential shared biological mechanisms underlying these interrelated diseases as well as the pleiotropic effects of these KD genes. *COL4A2* is a critical component of the basement membrane, and loss of function leads to disordered capillary networks during angiogenesis^77,78^. The C-terminal portion of *COL4A2* is a potent inhibitor of angiogenesis, prevents proliferation and migration of endothelial cells and induces apoptosis^79^. Moreover, variants of *COL4A2* are implicated in vascular cell survival, atherosclerotic plaque stability and risk of myocardial infarction, as well as hemorrhagic stroke^80,81^.

The KD genes for HFpEF were mainly from the vascular smooth muscle contraction, phosphatidylinositol signal system and endocytosis pathways, which further highlight the functional roles of inflammation and systemic vascular remodeling in the pathogenesis of HFpEF. Two examples of genes intricately linked to vascular wall mechanics rather than cardiomyocyte mechanics include *MYH11* and *MYLK. MYH11* encodes one of the smooth muscle cell myosin heavy chains, and variants are associated with familial thoracic aneurysm syndrome^82,83^. *MYLK* encodes a myosin light chain kinase that is implicated in inflammatory responses, apoptosis, and vascular permeability. Variants of *MYLK* are associated with arterial and aortic aneurysmal disease^84,85^.

Several strengths and limitations need to be considered when interpreting these findings. First, this study represents the first attempt to systematically examine and integrate genetic variants for HF phenotypic subtypes using pathway and network approaches with special emphasis on revealing mechanistic similarities and differences between HFrEF and HFpEF with higher statistical efficiency. The second strength is the large, previously validated and high-quality phenotyping of women of different ethnic backgrounds, which allows the detection of HF mechanisms shared across ethnicities. Thirdly, two additional high-quality cohorts, JHS and FHS, served as replication populations supporting the robustness of our findings. One major limitation is that our results were based upon germline mutations. Therefore, it is unclear whether mutations in the identified genomic regions and pathways would impact downstream expression levels in particular tissues of interest, and whether the identified genes and pathways are up-/down-regulated before and after HF events. This highlights the critical need for future studies that will quantify the downstream gene expression changes by comparing population with and without HF. Moreover, it will be important to validate these results in men in order to examine the effect of sex on HF and to replicate our finding using suitable animal models of HF in order to further validate these newly discovered pathways.

## Conclusion

This study validated previously identified locus and defined novel loci for HF and its subtypes, implicating specific molecular pathways, some shared and others unique, that contribute to HF and its subtypes. We highlight the significant mechanistic role of the inflammation and vascular remodeling (angiogenesis and vessel patterning) in the genetic signals associated with HFpEF and HFrEF, supporting the concept that HF is largely a disease of the systemic vasculature. Finally, this work defines several leading and novel targets and pathways for risk stratification and design of potential new HF interventions.

## Acknowledgements

The WHI program is funded by the National Heart, Lung, and Blood Institute, National Institutes of Health, U.S. Department of Health and Human Services through contracts HHSN268201600018C, HHSN268201600001C, HHSN268201600002C, HHSN268201600003C, and HHSN268201600004C.

The Jackson Heart Study (JHS) is supported and conducted in collaboration with Jackson State University (HHSN268201800013I), Tougaloo College (HHSN268201800014I), the Mississippi State Department of Health (HHSN268201800015I) and the University of Mississippi Medical Center (HHSN268201800010I, HHSN268201800011I and HHSN268201800012I) contracts from the National Heart, Lung, and Blood Institute (NHLBI) and the National Institute on Minority Health and Health Disparities (NIMHD). The authors also wish to thank the staffs and participants of the JHS.

From the Framingham Heart Study of the National Heart Lung and Blood Institute of the National Institutes of Health and Boston University School of Medicine. This project has been funded in whole or in part with Federal funds from the National Heart, Lung, and Blood Institute, National Institutes of Health, Department of Health and Human Services, under Contract No. 75N92019D00031.

Dr. Alan R. Morrison is supported by the Research Project Grant NIH NHLBI R01HL139795, the Institutional Development Award (IDeA) from NIH NIGMS P20GM103652, and the Career Development Award Number 7IK2BX002527 from the United States Department of Veterans Affairs Biomedical Laboratory Research and Development Program.

## Disclaimer

The views expressed in this manuscript are those of the authors and do not necessarily represent the views of the National Heart, Lung, and Blood Institute; the National Institutes of Health, the Department of the Veterans Affairs; or the U.S. Department of Health and Human Services.

