## Supplements for "Biological Pathways and Gene Networks Link Inflammation and Vascular Remodeling to Both Heart Failure with Preserved and Reduced Ejection Fraction in Women across Ethnicities"

Qing Liu, Ph.D. ^1,2^, Kei Hang K. Chan, Ph.D. ^2,3^, Alan R. Morrison, M.D., Ph.D. ^4,5^, Stephen T. McGarvey, Ph.D. ^1^, Xi Luo, Ph.D. ^6^, James G. Wilson, M.D. ^7^, Adolfo Correa, M.D., Ph.D. ^8^, Alexander P. Reiner, M.D. ^9,10^, Jie Li, Ph.D. ^1,2^, *****Simin Liu, M.D., Sc.D. ^1,2,5^, *****Wen-Chih Wu, M.D. ^1,4,5^

^1^Department of Epidemiology, School of Public Health, Brown University, Providence, RI 02912, USA

^2^ Center for Global Cardiometabolic Health, Brown University, 121 South Main St, Providence, RI, 02903

^3^ Department of Biomedical Sciences, City University of Hong Kong, Hong Kong

^4^ Providence Veterans Affairs Medical Center, Providence, RI 02908, USA

^5^ Alpert Medical School at Brown University, Providence, RI 02903, USA

^6^ Department of Biostatistics and Data Science, School of Public Health, University of Texas Health Science Center at Houston, Houston, TX 77030, USA

^7^ Department of Physiology and Biophysics, University of Mississippi Medical Center, Jackson, MS 39216

^8^ Department of Medicine, University of Mississippi Medical Center, Jackson, MS 39216

^9^ Department of Epidemiology, University of Washington, Seattle, WA 98195, USA

^10^ Division of Public Health Sciences, Fred Hutchinson Cancer Research Center, Seattle, WA 98109, USA.

***Co-Corresponding Authors**:

Simin Liu

Center for Global Cardiometabolic Health and Department of Epidemiology, Brown University, 121 South Main St, Providence, RI, 02903

Wen-Chih Wu

Center of Innovation in Long Term Services & Support, Providence VA Medical Center, Providence, RI

Division of Cardiology, Department of Medicine, Alpert Medical School, Brown University, Providence, RI

### Supplemental Material

Supplemental Table 1. Common Name for Genes Close to Validated and Newly Discovered Heart Failure Loci.

|  | **Lead SNP** | **References** | **Position (hg19)** | **Close Gene ^a^** | **Common Gene Name** |
| --- | --- | --- | --- | --- | --- |
| **Validated loci** | | | | | |
|  | rs4420638 | He, et al^1^ | 19:45422946 | Near *APOE* | Apolipoprotein E |
|  |  |  |  | Near *APOC* | Apolipoprotein C |
| **Discovered loci** | | | | | |
|  | rs12719020 |  | 7:51066547 | Near *COBL* | Cordon-Bleu WH2 Repeat Protein |
|  | rs12067046 |  | 1:208913216 | Near *PLXNA2* | Plexin A2 |
|  | rs12599260 |  | 16:50093238 | Near *HEATR3* | HEAT Repeat Containing 3 |
|  | rs114553497 |  | 1:229507916 | Near *CCSAP* | Centriole, Cilia And Spindle Associated Protein |
|  |  |  |  | Near *ACTA1* | Actin, Alpha 1, Skeletal Muscle |
|  | rs10229703 |  | 7:134661230 | Near *CALD1* | Caldesmon 1 |
|  |  |  |  | Near *AGBL3* | ATP/GTP Binding Protein Like 3 |
|  | rs149663839 |  | 11:34408645 | Near *ABTB2* | Ankyrin Repeat And BTB Domain Containing 2 |
|  |  |  |  | Near *CAT* | Catalase |

^a^ Closest genes within ± 300 kb of the lead SNPs.

Supplemental Table 2. Pathways Identified Specifically for HFrEF by Multiple Pathway Methodologies ^a^.

| **Pathways** | **Discovery Findings ^b^** | |  | **Validation Findings ^c^** | | | | | | |
| --- | --- | --- | --- | --- | --- | --- | --- | --- | --- | --- |
|  | **HFrEF - EA** | **HFrEF - AA** |  | **HFrEF** | | |  | **HFpEF** | | |
|  |  |  |  | **EA** | **AA** | **HA** |  | **EA** | **AA** | **HA** |
| Vascular smooth muscle contraction | G, M |  |  | -- |  | G |  | G, M | G |  |
| Extracellular matrix-receptor interaction | G, M |  |  | -- | G | G |  | G, M | G | G |
| Cell adhesion molecules | G, M |  |  | -- | G | G |  | G | G, M | G, M |
| Intrinsic pathway | G, M |  |  | -- |  | M |  |  |  | M |
| Rac1 pathway | G, M |  |  | -- |  |  |  |  | M |  |
| Mucin type O-glycan biosynthesis |  | G, M |  |  | -- | M |  | G |  | G |
| NRAGE signals death through JNK |  | G, M |  |  | -- |  |  | G, M |  |  |
| Signaling by Rho GTPases |  | G, M |  |  | -- | G |  | M |  | G, M |
| Developmental biology |  | G, M |  | G | -- | G |  | G | G | M |
| Axon guidance |  | G, M |  | G | -- | G |  | G, M | G | G, M |
| Netrin-1 signaling |  | G, M |  | G | -- | G |  | G | G | G, M |

Abbreviations: AA (African American), EA (European American), JNK (JUN Kinase), HA (Hispanic American), HFpEF (heart failure with preserved ejection fraction), HFrEF (heart failure with reduced ejection fraction).

^a^ G represents GSA-SNP, M represents Mergeomics.

^b^ The discovery findings were significant pathways (FDR-adjusted q value < 0.2) for HFrEF identified from the Women’s Health Initiative Study (N=7,982 for AA and N=4,133 for EA) and validated from the Jackson Heart Study (N=1,853) and Framingham Heart Study (N= 1,755).

^c^ The validation findings were presented as pathways with nominal *P* value < 0.05 from the Women’s Health Initiative Study (N=7,982 for AA and N=4,133 for EA).

Supplemental Table 3. Pathways Identified Specifically for HFpEF by Multiple Pathway Methodologies ^a^.

| **Pathways** | **Discovery Findings ^b^** | |  | **Validation Findings ^c^** | | | | | | |
| --- | --- | --- | --- | --- | --- | --- | --- | --- | --- | --- |
|  | **HFpEF - EA** | **HFpEF - AA** |  | **HFrEF** | | |  | **HFpEF** | | |
|  |  |  |  | **EA** | **AA** | **HA** |  | **EA** | **AA** | **HA** |
| Axon guidance | G, M |  |  | G | G, M | G |  | -- | G | G, M |
| Phosphatidylinositol signal system | G, M |  |  |  |  |  |  | -- | G, M | G, M |
| Adherens junction | G, M |  |  |  | M | G |  | -- | G | G |
| Pre-NOTCH expression and processing | G, M |  |  |  |  |  |  | -- |  | M |
| HS-GAG degradation | G, M | G, M |  |  |  |  |  | -- | -- |  |
| HS-GAG biosynthesis | G, M |  |  |  | G |  |  | -- | G, M |  |
| Ion transport by P-type ATPases | G, M |  |  |  | G, M |  |  | -- |  | G, M |
| Ion channel transport | G, M |  |  |  |  | G |  | -- |  | G |
| Cell adhesion molecules |  | G, M |  | G, M | G | G |  | G | -- | G, M |
| Long-term depression |  | G, M |  | G |  | G |  |  | -- |  |
| Endocytosis |  | G, M |  |  | M | G, M |  | G | -- | G, M |
| Signaling by BMP |  | G, M |  |  |  |  |  |  | -- |  |
| The role of Nef in HIV-1 replication and disease pathogenesis |  | G, M |  |  | M |  |  |  | -- | M |
| Cell-cell junction organization |  | G, M |  |  |  | G |  |  | -- |  |

Abbreviations: AA (African American), BMP (bone morphogenetic protein), EA (European American), GAG (glycosaminoglycan), HA (Hispanic American), HFpEF (heart failure with preserved ejection fraction), HS (heparan sulfate).

^a^ G represents GSA-SNP, M represents Mergeomics.

^b^ The discovery findings were significant pathways (FDR-adjusted q value < 0.2) for HFpEF identified from the Women’s Health Initiative Study (N=7,982 for AA and N=4,133 for EA) and validated from the Jackson Heart Study (N=1,853) and Framingham Heart Study (N= 1,755). Pathways shared with HFrEF were not included.

^c^ The validation findings were presented as pathways with nominal *P* value < 0.05 from the Women’s Health Initiative Study (N=7,982 for AA and N=4,133 for EA).

Supplemental Table 4. Characteristics of Identified Significant Pathways for Heart Failure by Ethnicities

|  | **Pathways** | **No. of genes** | **Min. *P* value** | **Main functions** |
| --- | --- | --- | --- | --- |
| **European Americans – HFrEF (n=5)** | | | | |
|  | Vascular smooth muscle contraction (VSMC) | 115 | 4.29×10^-4^ | VSMC is a highly specialized cell whose principal function is contraction. On contraction, VSMCs shorten, thereby decreasing the diameter of a blood vessel to regulate the blood flow and pressure^2^. The dysfunction of VSMC is directly related to hypertension, a well-understood risk factor of HF^3^. Meanwhile, many other mechanisms, especially inflammation and subsequent reduction-oxidation reaction, are also involved in regulating vascular smooth muscle function. Impaired endothelial and vascular smooth muscle dysfunction are associated with a poor prognosis in HF^4^. |
|  | Extracellular matrix (ECM)-receptor interaction | 84 | 3.79×10^-5^ | The ECM consists of a complex mixture of structural and functional macromolecules and serves an important role in tissue and organ morphogenesis and in the maintenance of cell and tissue structure and function^2^. ECM contributes to the angiogenesis and vascular patterning via multiple ways. ECM interacts with vascular endothelial growth factor (VEGF)-A, modulating its availability, its gradient organization and its signaling properties. ECM can indirectly signal through the Notch pathway by engaging a specific array of integrins. Moreover, ECM influences cellular tension and cytoskeleton organization, which are key aspects during sprouting^5^. In addition, the cardiac ECM (primarily collagen I) provides a platform for cardiomyocytes to maintain structure and function, and the change in ECM properties following an insult strongly drives the progression toward HF via multiple mechanisms, including myocardial fibrosis and changed ECM protein orientation^6,7^. |
|  | Cell adhesion molecules (CAMs) | 134 | 3.32×10^-6^ | CAMs are glycoproteins expressed on the cell surface and play a critical role in a wide array of biologic processes, including hemostasis, immune response, inflammation, etc.^2^ The CAMs are also critical players in the angiogenic cascade. For example, CAMs interact with endothelial cells by modulating the processes of cell adhesion, migration and proliferation, and with ECM molecules in the response to VEGF^8^. In addition, the CAMs play functional roles in a wide range of other biological processes including homeostasis, immune response, and inflammation^9^, and had been connected with metabolic syndrome^10^ and coronary heart disease^11^ in different ethnic populations. |
|  | Intrinsic prothrombin activation pathway | 23 | 2.85×10^-5^ | The second phase of blood coagulation or clotting – the activation of prothrombin^12^. |
|  | Rac-1 pathway | 23 | 2.68×10^-38^ | Rac-1 is a small G-protein in the Rho family that regulates cell motility in response to extracellular signals^12^. |
| **African Americans – HFrEF (n=6)** | | | | |
|  | Mucin type O-glycan biosynthesis | 30 | 6.42×10^-5^ | O-glycans are a class of glycans that modify serine or threonine residues of proteins. Mucins are highly O-glycosylated glycoproteins ubiquitous in mucous secretions on cell surfaces and in body fluids^2^. |
|  | NRAGE signals death through JUN Kinase (JNK) | 43 | 2.28×10^-5^ | Once bound by either NGF or proNGF, p75NTR interacts with NRAGE, thus leading to phosphorylation and activation of JNK. JNK controls apoptosis in two ways: it induces transcription of pro-apoptotic genes, and directly activates the cell death machinery^13^. |
|  | Signaling by Rho GTPases | 113 | 1.04×10^-7^ | Rho GTPases belong to the Rho family, typically binary switches controlling a variety of biological processes, including dynamic rearrangements of the plasma membrane-associated actin cytoskeleton, regulating actomyosin contractility and microtubule dynamics, cell growth control, cytokinesis, cell motility, cell-cell and cell-extracellular matrix adhesion, cell transformation and invasion, and development^13^. |
|  | Developmental biology | 396 | 6.18×10^-5^ | This pathway includes processes that a fertilized egg gives rise to the diverse tissues of the body^13^. |
|  | Axon guidance | 251 | 7.64×10^-8^ | This pathway is the process by which neurons send out axons to reach the correct targets^13^. Although the main function of axon guidance pathway is related to localization and neuronal extension, molecules within the pathway had been connected to angiogenesis and vascular patterning. For instance, axon guidance molecules and receptors, including semaphorins (SEMAs), plexins (PLXNs) and neuropilins (NRPs), slits and roundabouts (ROBOs), and netrins (NTNs) and netrin receptors (UNC5A-D), play functional roles in vascular development^14^. The molecules can either be directly linked to sprouting and guidance of capillary tip cells for angiogenesis, or have involved to vasculature, especially modulation of the activity of the VEGF signaling pathway, thus affecting guided vascular patterning by rendering vessels more or less responsive to VEGF^14^. |
|  | Netrin-1 signaling | 41 | 2.38×10^-5^ | Netrins are proteins that play a crucial role in neuronal migration and in axon guidance. Netrin-1 is the most studied member of the family and plays a crucial role in neuronal navigation during nervous system development mainly through its interaction with its receptors DCC and UNC5^13^. |
| **European Americans – HFpEF (n=8)** | | | | |
|  | Phosphatidylinositol signal system | 76 | 1.63×10^-5^ | Phosphatidylinositol is a small lipid molecule which can be phosphorylated by a host of lipid kinases to produce a variety of phosphatidylinositol monophosphates (PI3P, PI4P, and PI5P), diphosphates, and a triphosphate that are collectively known as phosphoinositides^2^. Phosphatidylinositol signaling system regulates a broad of molecular functions, which involve vascular patterning, inflammation, and general cell signaling. For example, phosphoinositide 3-kinase (PI3K) signaling mediates angiogenesis and expression of VEGF in endothelial cells^15^, and it regulates cardiomyocyte size, survival and inflammation in cardiac hypertrophy and HF, probably by interacting with calcium signaling^16,17^. |
|  | Adherens junction (AJ) | 75 | 1.19×10^-9^ | Cell-cell AJ is the most common type of intercellular adhesions, and is important for maintaining tissue architecture and cell polarity and can limit cell movement and proliferation^2^. AJs are ubiquitous along the vascular tree. AJs in endothelial cells regulate angiogenesis and vessel maintenance. For example, vascular endothelial cadherin and its intracellular partners mediate adhesion via multiple mechanisms, including the direct activation of phosphatidylinositol -3 (PI3) kinase and Rac, and the formation of complexes with VEGF receptors^18^. Fascia AJ is one category of intercalated disks which maintain structural integrity and synchronized contraction of cardiac tissue. One component of this AJ, β-catenin, had been linked to hypertrophic cardiomyopathy and end-stage HF, suggesting some AJ proteins may have unique functions in different types or at least in different stages of HF^19^. |
|  | Pre-NOTCH expression and processing | 44 | 2.45×10^-6^ | This pathway is the expression and processing of nascent forms of NOTCH precursors, which undergo extensive posttranslational modifications in the endoplasmic reticulum and Golgi apparatus to become functional^13^. |
|  | Heparan sulfate (HS)- glycosaminoglycan (GAG) degradation | 20 | 8.81×10^-5^ | This pathway is the degradation of heparan sulfate/heparin. HS-GAG, once combined with the core proteins to generate HS proteoglycans (HSPGs)^20^, can interact with growth factors, lipoproteins, cytokines or other proteins in the progression of various disorders, including atherosclerosis and diabetes^13^. HSPGs had been observed to involve in angiogenesis by mediating the angiogenic activity of the VEGF receptor 2^21^. HSPGs are also found to interact with lipoproteins, cytokines or other proteins in the progression of atherosclerosis and diabetes^22^. An animal study also found HS-GAG functions might be associated with aging, thus partly supporting HFpEF as an age-related disorder^23^. |
|  | HS-GAG biosynthesis | 31 | 3.73×10^-4^ | This pathway is the biosynthesis of heparan sulfate/heparin^13^. |
|  | Axon guidance | 251 | 1.44×10^-9^ | Also found for HFrEF among African Americans. |
|  | Ion transport by P-type ATPases | 34 | 2.92×10^-6^ | The P-type ATPases are a large group of ion and lipid pumps, and serve as a basis for nerve impulses, relaxation of muscles, secretion and absorption in the kidney, absorption of nutrient in the intestine and other physiological processes^13^. |
|  | Ion channel transport | 55 | 8.45×10^-6^ | This is a group of ion channels that mediate the flow of ions across the plasma membrane of cells^13^. |
| **African Americans – HFpEF (n=7)** | | | | |
|  | Cell adhesion molecules | 134 | 6.88×10^-6^ | Also found for HFrEF among European Americans. |
|  | Long term depression | 70 | 1.14×10^-5^ | It is a process involving a decrease in the synaptic strength between parallel fiber and Purkinje cells, and it is a molecular and cellular basis for cerebellar learning^2^. |
|  | Endocytosis | 183 | 2.76×10^-4^ | Endocytosis is a mechanism for cells to remove ligands, nutrients, and plasma membrane proteins, and lipids from the cell surface, bringing them into the cell interior^2^. Endocytosis involved in a variety of cell functions which can be linked to angiogenesis and cardiac dysfunction. For example, endocytosis regulates VEGF signaling during angiogenesis through spatial control of VEGF receptors endocytosis^24^, and also involves in the desensitization and internalization of β-adrenergic receptors (βARs), which are critical regulators of cardiovascular physiology^25^. Importantly, PI3K, as previously described, has also been found in regulating βARs endocytosis^26^. In addition, a group of endocytic regulatory proteins – C-terminal Eps15 homology domain-containing proteins (EHDs) had been linked to Na^+^ and Ca^2+^ homeostasis, which is fundamental for cardiac arrhythmia^27^. |
|  | Signaling by Bone morphogenetic proteins (BMP) | 23 | 2.93×10^-28^ | BMPs are members of the Transforming growth factor-Beta (TGFB) family. BMP signaling is linked to a wide variety of clinical disorders, including vascular diseases, skeletal diseases and cancer^13^. |
|  | The role of Nef in HIV-1 replication and disease pathogenesis | 28 | 1.24×10^-9^ | This pathway plays an important role in several steps of HIV replication^13^. |
|  | HS-GAG degradation | 20 | 9.34×10^-7^ | Also found for HFpEF among European Americans. |
|  | Cell-cell junction organization | 56 | 7.77×10^-8^ | Epithelial cell-cell contacts consist of three major adhesion systems: adherens junctions (AJs), tight junctions (TJs), and desmosomes. These adhesion systems have different functions and compositions. AJs play a critical role in initiating cell-cell contacts and promoting the maturation and maintenance of the contacts. TJs form physical barriers in various tissues and regulate paracellular transport of water, ions, and small water-soluble molecules. Desmosomes mediate strong cell adhesion linking the intermediate filament cytoskeletons between cells and playing roles in wound repair, tissue morphogenesis, and cell signaling^13^. |

Abbreviations: HF (heart failure), HFpEF (heart failure with preserved ejection fraction), HFrEF (heart failure with reduced ejection fraction).

A. B.

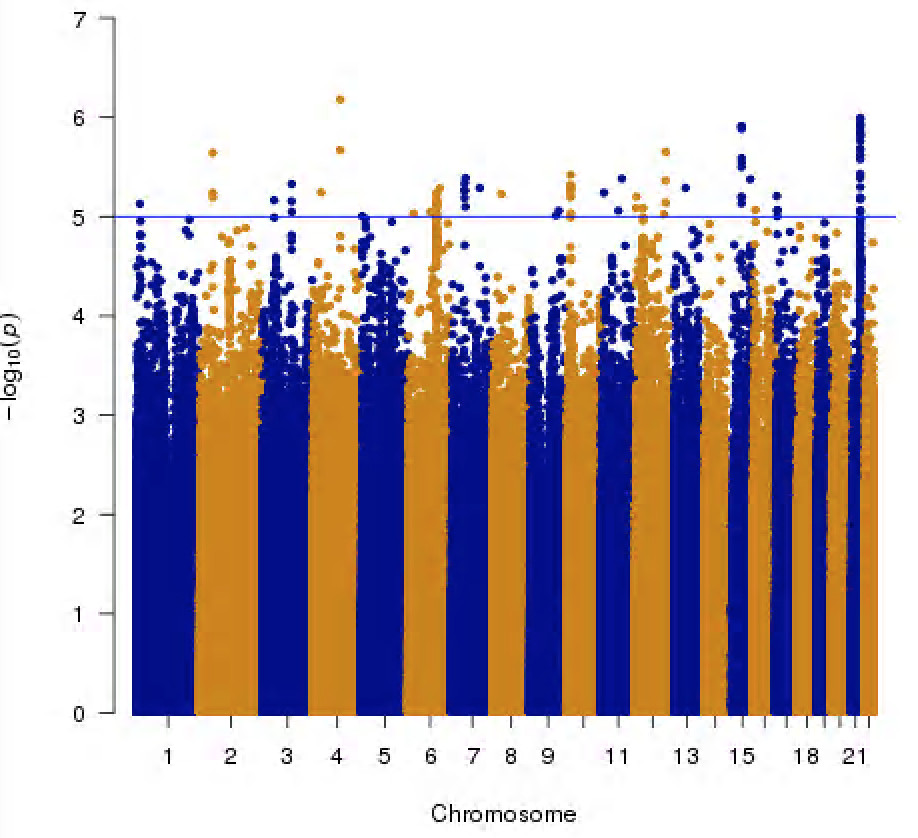

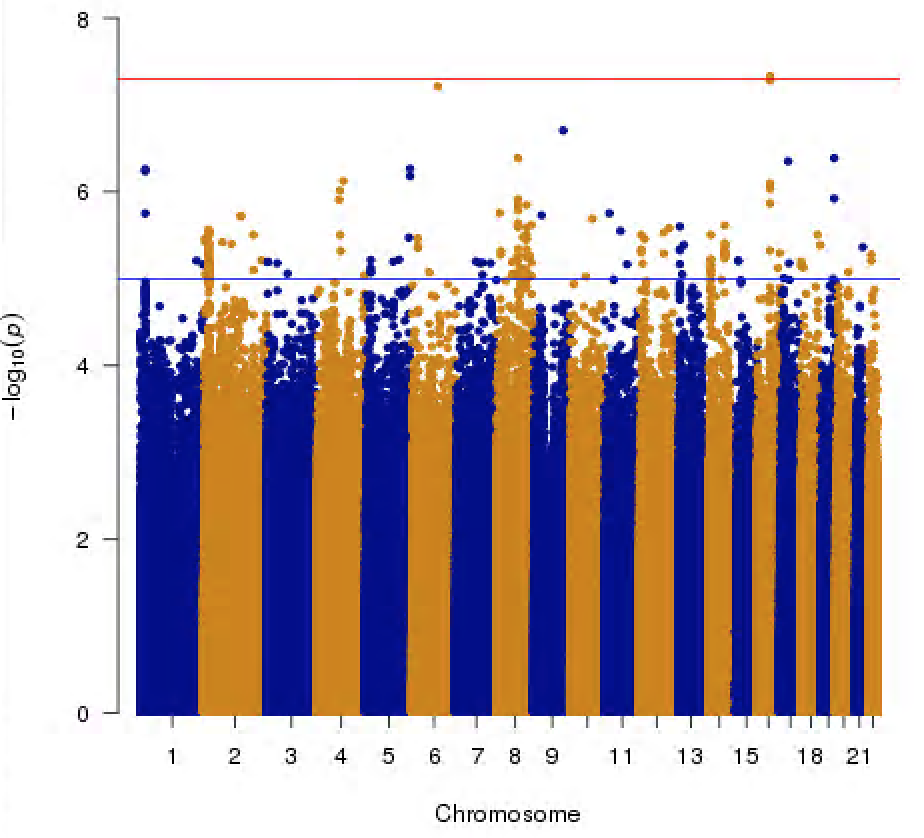

C. D.

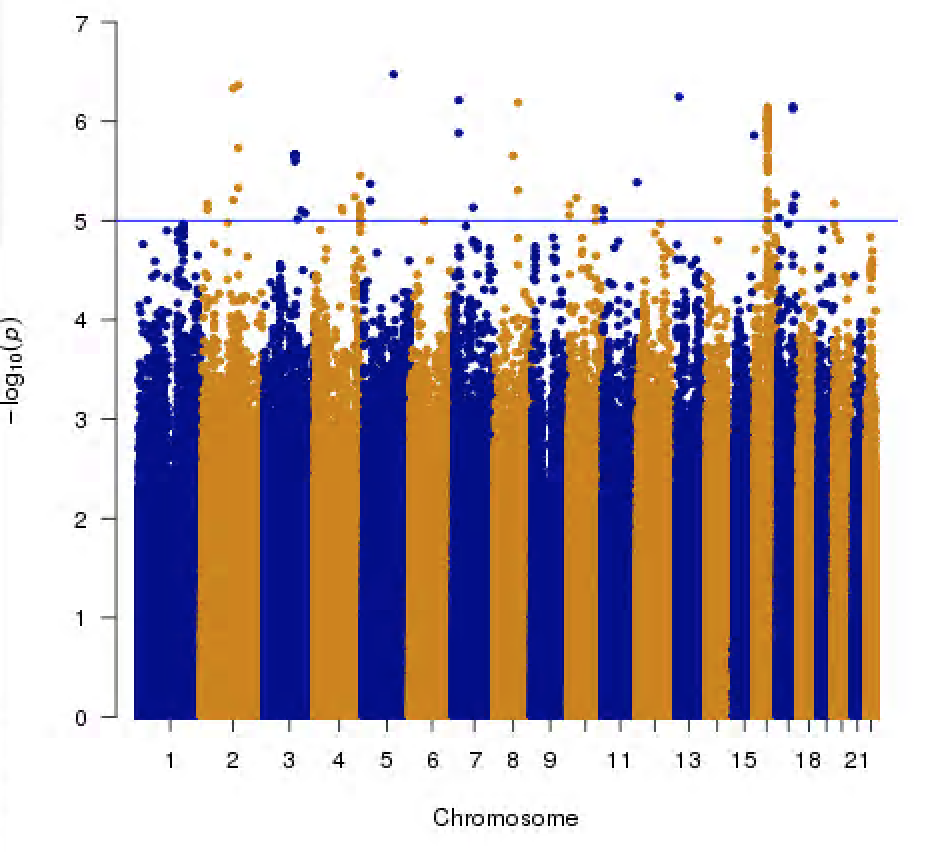

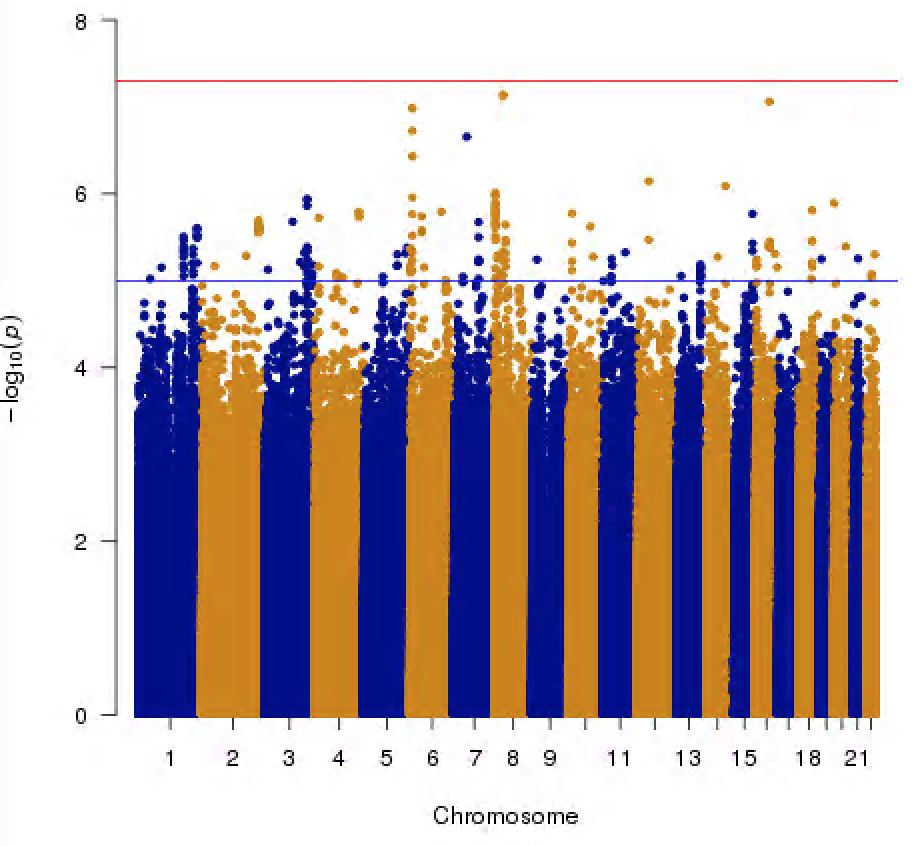

Supplemental Figure 1. Manhattan plots of heart failure among African Americans (n=7,982) and European Americans (n=4,133) in the Women’s Health Initiative Study.

(A: HFrEF among WHI-EA; B: HFrEF among WHI-AA; C: HFpEF among WHI-EA; D: HFpEF among WHI-AA)

A.

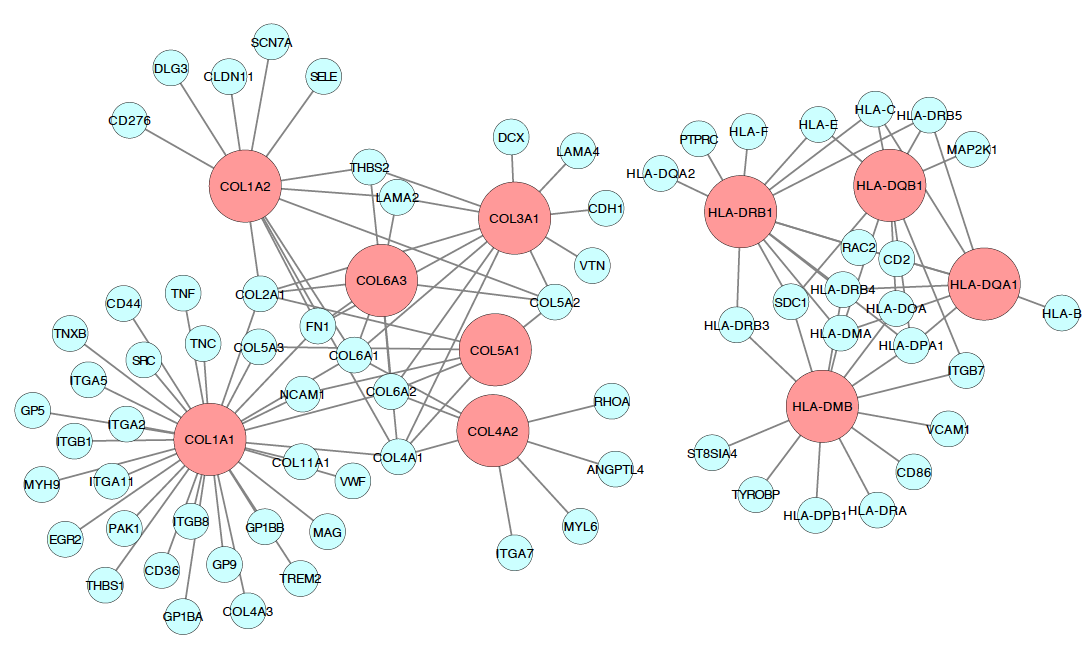

B.

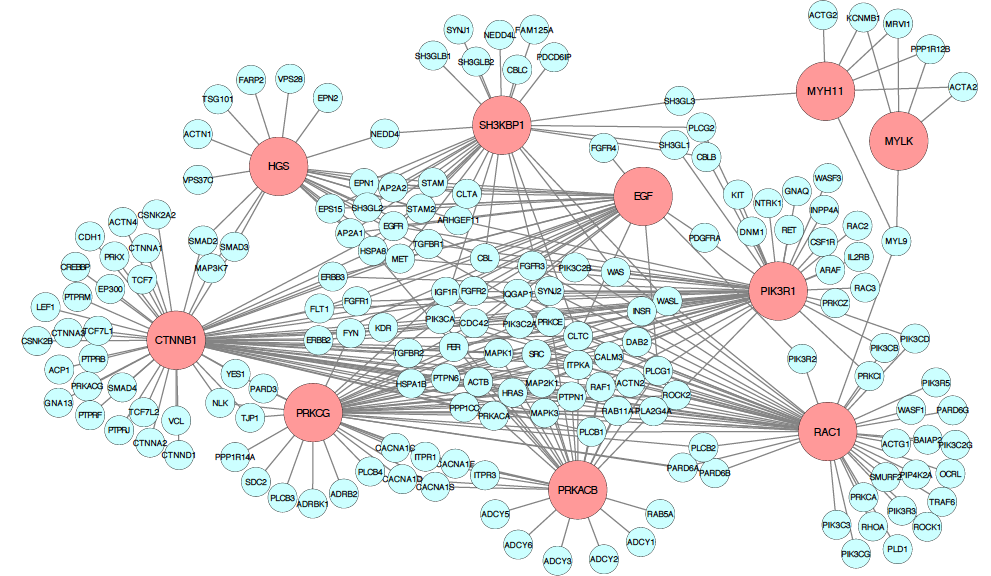

Supplemental Figure 2. Network key driver genes of the pathways enriched for HFrEF and HFpEF among African and European American Women.

(A: HFrEF and HFpEF shared; B: HFpEF specific)

Top 10 ranked multi-tissue key driver genes of the heart failure pathways in the Bayesian network and the protein-protein interaction network. The modes represent genes, and the edge shows the interaction. The color nodes are: light red, top 10 ranked key driver genes; mint, genes interact with the key drivers. The figure was created using Cytoscape^28^.

21.	Chiodelli P, Mitola S, Ravelli C, Oreste P, Rusnati M, Presta M. Heparan Sulfate Proteoglycans Mediate the Angiogenic

Activity of the Vascular Endothelial Growth Factor

Receptor-2 Agonist Gremlin. Arterioscler Thromb Vasc Biol 2011;31:e116-e27.
